# Comparative analysis highlights variable genome content of wheat rusts and divergence of the mating loci

**DOI:** 10.1101/060665

**Authors:** Christina A. Cuomo, Guus Bakkeren, Hala Badr Khalil, Vinay Panwar, David Joly, Rob Linning, Sharadha Sakthikumar, Xiao Song, Xian Adiconis, Lin Fan, Jonathan M. Goldberg, Joshua Z. Levin, Sarah Young, Qiandong Zeng, Yehoshua Anikster, Myron Bruce, Meinan Wang, Chuntao Yin, Brent McCallum, Les J. Szabo, Scot Hulbert, Xianming Chen, John P. Fellers

## Abstract

Three members of the *Puccinia* genus, *P. triticina* (*Pt*), *P. striiformis f.sp. tritici*(*Pst*), and *P. graminis f.sp. tritici* (*Pgt*), cause the most common and often most significant foliar diseases of wheat. While similar in biology and life cycle, each species is uniquely adapted and specialized. The genomes of *Pt* and *Pst* were sequenced and compared to that of *Pgt* to identify common and distinguishing gene content, to determine gene variation among wheat rust pathogens, other rust fungi and basidiomycetes, and to identify genes of significance for infection. *Pt* had the largest genome of the three, estimated at 135 Mb with expansion due to mobile elements and repeats encompassing 50.9% of contig bases; by comparison repeats occupy 31.5% for *Pst* and 36.5% for *Pgt*. We find all three genomes are highly heterozygous, with *Pst* (5.97 SNPs/kb) nearly twice the level detected in *Pt* (2.57 SNPs/kb) and that previously reported for *Pgt*. Of 1,358 predicted effectors in *Pt*, 784 were found expressed across diverse life cycle stages including the sexual stage. Comparison to related fungi highlighted the expansion of gene families involved in transcriptional regulation and nucleotide binding, protein modification, and carbohydrate enzyme degradation. Two allelic homeodomain, HD1 and HD2, pairs and three pheromone receptor (*STE3*) mating-type genes were identified in each dikaryotic *Puccinia* species. The HD proteins were active in a heterologous *Ustilago maydis* mating assay and host induced gene silencing of the HD and *STE3* alleles reduced wheat host infection.

## Introduction

Rust fungi have the most complex life cycles among described fungi, with many stages having discrete morphologies and very distinguishable sexual and asexual propagation. For many rust fungi, these stages occur on two different, unrelated host plants requiring two different infection strategies (heteroecious). For many, such as the cereal rust fungi, the asexual stage can successfully propagate and lead to epidemics as long as the host is present. When biologically stressed, the fungus enters the sexual cycle and survival teliospore structures are produced. In homoecious rust fungi, like flax rust (*Melampsora lini* (Ehrenb.) Lév.), both stages evolved on the same host (autoecious). In heteroecious rust fungi, such as the cereal rust pathogens, major host jumps occur through evolution of the asexual (uredinial) stage to infect a different host (Savile 1976). These complex interactions result in the production of up to five different rust spore types, requiring very discrete developmental programs, resulting in altered gene expression profiles.

The obligate biotrophic lifestyle of wheat rust pathogens hampers the ability to culture the fungus *in vitro* and thus limits biological studies. Genetic studies by crossing individual strains is not trivial due to the difficulty of breaking teliospore dormancy in order to infect the alternate hosts. Most of what is known about wheat rust pathogen biology is based on extensive cytology (Bushnell and Roelfs 1984) and isolate interactions with host resistance genes. Many interactions between rust fungi and their cereal hosts have been shown to genetically conform to the gene-for-gene theory (Flor 1942; Loegering, and Powers 1962). The majority of wheat rust resistance genes (McIntosh *et al*. 1995) have been shown to be dominant or semi-dominant (Statler 1979, 1982, 2000) and current models imply an interaction between the resistance genes and fungal effectors (Sperschneider *et al*. 2013; Petre *et al*. 2014).

Wheat leaf rust, caused by *Puccinia triticina* Eriks (*Pt*), is the most commonly occurring cereal rust disease worldwide. Leaf rust on wheat was first recognized as different from stem rust in 1718 (De Candolle 1815) and included into a species complex (*P. rubigo-vera*) and taxonomic refinements resulted in the current classification based on differences in spore morphology and alternate host range (Eriksson 1899; Savile 1984; Anikster *et al*. 1997). *Pt* is an obligate biotroph that can complete its sexual cycle on either of two known alternate host species, *Thalictrum speciosissimum* Loefl. (meadow rue; (Jackson and Mains 1921; Saari *et al*. 1968)) or *Isopyrum* (Brizgalova 1935, 1937). The complete *Pt* cycle consists of five spore stages (Bolton *et al*. 2008). The urediniospore is the most common and is asexual and polycyclic. At maturity when leaves begin to senesce, the fungus will form black teliospores as survival structures on the underside of the leaf. Within the teliospores, karyogamy takes place and promycelia are formed when the teliospores germinate. Four haploid basidiospores, in which the mating types have segregated, are formed and infect the alternate host, forming the pycnium. Dikaryotization occurs through fusion between a receptive flexous hyphae and a pycniospore of a different mating type. After fusion, the dikaryotic state is reestablished and an aecium will form on the underside of the leaf from which aeciospores will be released and travel to the wheat host. After landing on wheat leaves, the aeciospore will germinate forming an appressorium over a stoma. The haustorial mother cell forms in the substomatal cavity and attaches to the host cell wall. The plant cell wall is breached between 24 to 48 hours, forming haustoria. The fungus will spread intercellularly and a uredinium is formed at 7 days, from which urediniospores are produced to complete the life cycle. Wheat suffers from two other major rust diseases: stem rust, caused by *Puccinia graminis* Pers.:Pers. f. sp. *tritici* Erikss. & E. Henn. (Pgt; (Leonard and Szabo 2005) and stripe rust, caused by *P. striiformis* Westend. f. sp. *tritici* Erikss. (*Pst*; (Chen *et al*. 2014). While all three wheat rust pathogens share very similar biology and spore stages, only *Pgt* and *Pst* have *Berberis* spp. as an alternate host (Statler 1979, 1982, 2000; Jin *et al*. 2010).

Rust fungi belong to the subphylum Pucciniomycotina that together with the Ustilaginomycotina, the true smut fungi, and the Agaricomycotina, which include mushroom-forming species, make up the phylum Basidiomycota. In this subphylum, the sexual cycle typically requires cell-cell fusion governed by both pheromone (*mfa*) and pheromone receptor (*STE3*) genes, which then allows the formation of heterodimeric transcription factors coded for by two classes of homeodomain-containing protein genes, *HD1* and *HD2* (Raudaskoski and Kothe 2010; Kues *et al*. 2011). In the corn smut fungus *Ustilago maydis*, the mating-type locus contains both the pheromone receptor gene *Pra* (the *STE3* equivalent) and a pheromone precursor gene *mfa* (Brefort *et al*. 2009). In all basidiomycetes studied to date, heterodimeric HD-containing transcription factors have been implicated in the mating process. They are found in pairs of genes each encoding subunits of a HD1 and HD2-containing protein that are divergently transcribed. Originally found with their start sites within 1 kb in *Um*, many variations exist and multiple pairs in arrays of linked diverged copies are often found in mushrooms (Casselton and Kües 2007; Raudaskoski and Kothe 2010) though single gene pairs are predicted in *Pleurotus djamor* (James *et al*. 2004) and *Pholiota nameko* (Yi *et al*. 2009). To date, the mating loci of the wheat rusts have not been carefully analysed.

As for other obligate pathogens, genome sequencing of rust fungi has advanced the basic understanding of these organisms, which are otherwise recalcitrant to laboratory study. Initial molecular analyses and phylogenetic data indicated that within each lineage of these three rust pathogens, adaptation to the wheat host had occurred independently (Zambino and Szabo 1993). Genome differences were identified in EST sequencing studies where it was shown that 40% of *Pt* ESTs did not have a match to *Pgt* and *Pst* (Xu *et al*. 2011). Comparison of contigs from BAC and genome sequencing have shown synteny between the three genomes, however, there are regions of gene insertions, expansion by mobile elements, and inversions (Cantu *et al*. 2011; Fellers *et al*. 2013).

Prior to this work, the genomes of *Pgt* and the poplar leaf rust *M. larici-populina (Mlp*) were sequenced and compared. Out of 17,773 and 16,399 predicted genes, respectively, a core set of genes was identified representing the biotrophic nature of rusts (Duplessis *et al*. 2011). More recently, a second *de novo* genome assembly of *Pgt* was completed (Upadhyaya *et al*. 2014). Three genome sequencing projects have been described for *Pst* (Cantu *et al*. 2011, 2013; Zheng *et al*. 2013). The first two focused on identifying the effector complement, and the third study on identifying heterozygosity between two isolates of *Pst*. In each study, the total number of predicted genes varied across the projects (22,815 versus 25,288, respectively). The genome of the flax rust *M. lini* (Ml/), has also been sequenced (Nemri *et al*. 2014). Comparative analyses with other basidiomycete genomic resources have provided initial insights into relatedness of subsets of genes (Xu *et al*. 2011; Nemri *et al*. 2014) but a comprehensive analysis among wheat rusts is missing.

Here, we have generated draft genome sequences of the wheat rust fungi *Pt* race 1 (BBBD) and *Pst* race PST-78 and updated the gene set of *Pgt* race SCCL, and utilized these sets to define the shared and unique properties of these three related pathogens. To examine gene content evolution, we compared predicted proteins to those of other high-quality basidiomycete genomes. We examined the three wheat rust pathogens for conservation of gene families, including effector genes, and compared to other sequenced rust fungi. We identified predicted secreted proteins including gene families found only in the wheat rust pathogens and found differences in expression levels between *Pt* life cycle stages, including sexual stages on the alternate host and wheat infection. In addition, we analyzed the mating-type gene complexes, revealed their evolutionary placement among basidiomycetes, tested the functionality of several *Pt* homeodomain proteins through heterologous expression in *U. maydis* (*Um*), and demonstrated a role for mating-type genes during wheat infection by host-induced gene silencing (HIGS).

## MATERIAL AND METHODS

For detailed descriptions of isolates, sequencing strategies, genome assemblies and annotation, polymorphism analyses, DNA and RNA isolation procedures, RNA sequencing, and cloning methods, see Supplementary Materials and Methods.

### *Puccinia* isolates and growth conditions

*Pt* race 1, BBBD was selected as the race to be sequenced. This isolate was first collected in 1954 (Ordoñez and Kolmer 2009) and represents the earliest race characterized into North America. For *Pst*, isolate 2K41-Yr9 was selected (race PST-78), which was collected from the Great Plains in 2000. PST-78 is a representative of races that were first identified in the U.S. in 2000 and then subsequently identified in other countries (Wellings *et al*. 2003; Hovmøller *et al*. 2008).

### DNA and RNA isolation

Genomic DNA was isolated from *Pt* and *Pst* urediniospores. RNA was isolated from three stages of *Pt* race 1: from fresh mature, “dormant” urediniospores that had been collected at 10 days post inoculation (DPI), from urediniospores that were germinated on water, but harvested at 8 hrs post germination initiation, and thirdly, from heavily infected wheat tissue at 6 DPI to represent the formation of urediniospores, initiation of secondary infection, and most of the infection structures. RNA was also isolated and sequenced for two stages for both *Pt* and *Pgt* from the alternate host, *Thalictrum speciosissimum* (meadow rue), and represented pycnia with their pycniospores and a mixture of pycnia and aecia with aeciospores. RNA was isolated from two stages of *Pst*, infected wheat tissue at 8 DPI and isolated haustoria, purified as described (Yin *et al*. 2009).

### Genome sequencing and assembly

For *Pt* genome sequencing, various sizes of genomic DNA libraries and platforms were used. In short, libraries of 3 kb and 8 kb fragment inserts were sequenced using Roche 454 FLX chemistry and two large insert libraries were end-sequenced using Sanger technology: a 40 kb insert Fosmid library (30,731 clones) and ~100 kb insert BAC library (15,000 clones (Fellers *et al*. 2013); Table S1). An initial assembly of FLX and Sanger data was generated with Arachne (HybridAssemble) (Jaffe *et al*. 2003). The assembly was updated to incorporate the FLX+ data by first generating a new *de novo* assembly of all data using Newbler runAssembly, with parameters-het and-large, and merging the output with contigs uniquely present in the first assembly.

Three similar *Pst* genomic DNA insert libraries were sequenced using FLX chemistry with a Roche 454. In addition, paired end Illumina reads were generated for three additional library sizes: fragment, 3–5 kb insert, and 40 kb Illumina-adapted Fosmids (Fosill library, (Williams *et al*. 2012); Table S2). Three initial assemblies were generated using different algorithms: Life Technologies’ Newbler program, the CLC (Qiagen, Hilden, Germany) denovo assembler, and ALLPATHS-LG (Gnerre *et al*. 2011). To provide the most complete representation of the genome, contigs from the ALLPATHS-LG assembly were first selected, and then unique contigs from the CLC assembly were incorporated; see Supplementary Methods for details.

### Polymorphism analysis

Heterozygous positions within the sequenced isolates of *Pt* and *Pst* were identified from Illumina data. Reads were aligned to each assembly using BWA (v0.5.9) (Li and Durbin 2010), and SNP positions were identified with GATK v2.1.9 UnifiedGenotyper, and then filtered by GATK VariantFiltration; see Supplementary Methods for details.

### RNA sequencing

Strand-specific libraries were constructed with poly(A) selected RNA samples using the dUTP second strand marking method (Parkhomchuk *et al*. 2009; Levin *et al*. 2010) for most samples. See Supplementary Methods for details of library construction and expression analysis.

### Genome annotation

Gene sets were annotated by incorporating RNA-Seq data and predicted gene structures from multiple de novo predictions as previously described (Haas *et al*. 2011); see supplementary information for detailed description.

### Cloning, expression, and functional analysis of *Pt* mating-type genes

The various *Pt* HD mating-type alleles were amplified by PCR from cDNA generated from total RNA isolated at 5 DPI from infected wheat cv. Thatcher leaves infected with *Pt* race 1 or from urediniospores germinated over water. These alleles were subsequently cloned in a *Ustilago-specific*, integrative vector for heterologous expression from the strong constitutive Hsp70 promoter and terminator elements. Constructs were stably transformed in to Um518 (Kronstad and Leong 1989) or strain FB1 (Banuett 1991), and in *U. hordei*. To test the function of various mating-type genes during infection of wheat by *Pt*, HIGS experiments were performed. To create the RNAi silencing vectors, fragments of size 393-bp, 430-bp, 351-bp and 345-bp of the genes *PtbWI, PtbEI, PtSTE3.3* and *PtSTE3.1*, respectively, were amplified by PCR, cloned into the vector pENTR/D-TOPO (ThermoFisher), and subsequently recombined with the binary destination vector pIPK007 (Himmelbach *et al*. 2007) using the LR GateWay recombination reaction to create the silencing vectors pRNAi-*PtbW1*, pRNAi-*PtbE1*, pRNAi-*PtSTE3.3* and pRNAi-*PtSTE3.1*, respectively. Using *Agrobacterium tumefaciens* strain C0R308, agroinfiltration assays, subsequent challenge by *Pt* and fungal biomass measurements using quantitative PCR measurements were performed as described previously (Panwar *et al*. 2013). For details on these procedures, see Supplementary Materials and Methods.

## RESULTS

### Genome expansion in *Pt* associated with repetitive element proliferation

High quality genome assemblies of *Pt* and *Pst* were generated by combining data from multiple sequencing technologies. A range of insert size libraries for both genomes were sequenced using Roche 454, Illumina and Sanger technologies (Tables S1 and S2). The assembled genome of *Pt* was the largest of the three wheat rust pathogens, totaling 135.3 Mb (Table 1); this assembly included 14,820 scaffolds of an N50 length of 544 Kb. The assembly of *Pst* totaled 117.31 Mb; this is comparable to previously reported values (Cantu *et al*. 2011; Zheng *et al*. 2013). The *Pst* assembly consisted of 9,715 scaffolds with N50 length of 519 Kb. RNA-Seq was used to guide gene prediction for both *Pt* and *Pst*, and to improve the gene set of *Pgt* (Methods). Of the three rust pathogens, *Pst* had the highest number of genes predicted with 19,542, though fewer than the number reported for other *Pst* genomes (Cantu *et al*. 2011; Zheng *et al*. 2013), while *Pt* had the smallest total of the three with 14,880 genes. All three genomes have high coverage of a core eukaryotic gene (CEG) set (Parra *et al*. 2007; Table 1). The CEG coverage of this *Pst* gene set (97%) is notably higher than that of the PST-130 assembly (66%), the only other publicly available *Pst* gene set, due to a higher fraction of partial gene matches in PST-130 (Figure S1). In addition, comparison of a larger set of basidiomycete conserved orthologs supports that few genes appear missing in the three wheat rust fungal genomes (Figure S2). Together, these gene conservation metrics suggest that these assemblies contain highly complete gene sets.

**Table 1.** Genome statistics for three wheat rusts, *Pt, Pst* and *Pgt*.

| Species | Assembly Size | % GC | Total Contig length | Scaffolds | Scaffold N50 | Contigs | Contig N50 | Genes | CEGMA (%)** |
| --- | --- | --- | --- | --- | --- | --- | --- | --- | --- |
| *P. triticina* BBBD, Race 1 | 135.34 Mb | 46.72 | 106.57 Mb | 14,820 | 544.26 Kb | 24,838 | 10.37 Kb | 14,880 | 240 (97) |
| *P. striiformis tritici* PST-78 | 117.31 Mb | 44.43 | 79.31 Mb | 9,715 | 519.86 Kb | 17,295 | 17.36 Kb | 19,542 | 240 (97) |
| *P. graminis tritici* * | 88.64 Mb | 42.35 | 81.52 Mb | 392 | 964.97 Kb | 4,557 | 39.5 Kb | 15,800* | 232 (94) |

\* updated from Duplessis *et al.* 2011, based on RNA-seq data.
\*\* CEGMA (248 Core Eukaryotic Gene Set) (Parra et al., 2007) match at $\geq 80\%$ alignment coverage.

The assemblies of the three wheat rust fungi varied significantly in size, ranging from 89 Mb for *Pgt* to 135 Mb for *Pt*. While the *Pst* assembly totals 117 Mb, the genome may in fact be smaller than the assembly size, as the high percentage of gaps (32%) in scaffolds suggests that small contigs fall in to some of the gap regions. By comparison, the *Pt* assembly consists of 21% and *the Pgt* assembly only 8% gap regions. While all assemblies are impacted by the high heterozygosity (see below), differences in repeat content and organization likely as well as the use of different sequencing technologies likely contribute to these differences. Each of the three wheat rust pathogen genomes was evaluated for content of repeated elements using both *de novo* predicted repeats and fungal elements from RepBase, which included 413 *Puccinia* sequences (Supplemental Material and Methods). The larger genome of *Pt* includes a higher fraction of repetitive elements, encompassing 50.9% of contig bases, whereas repeats covered only 31.5% of *Pst* and 36.5% of *Pgt* (Table 2). The expanded repeat content of *Pt* includes roughly 2-fold more of both Class I retroelements and Class II DNA elements. After excluding the identified repeats, the non-repetitive portions of the genomes are very similar, totaling 53.4 Mb for *Pt*, 54.4 Mb for *Pst*, and 51.8 Mb for *Pgt*.

**Table 2.** Repeat element content of the *Pt, Pst* and *Pgt* genomes

|  | *Pt* |  |  | *Pst* |  |  | *Pgt* |  |  |
| --- | --- | --- | --- | --- | --- | --- | --- | --- | --- |
|  | Elements (#) | Length (Mb) | Genome (%) | Elements (#) | Length (Mb) | Genome (%) | Elements (#) | Length (Mb) | Genome (%) |
| SINEs | 254 | 0.04 | 0.04 | 21 | <0.00 | 0.00 | 17 | <0.00 | 0.00 |
| LINEs | 1,380 | 0.51 | 0.48 | 344 | 0.26 | 0.33 | 419 | 0.32 | 0.40 |
| LTR elements | 27,819 | 17.16 | 16.10 | 9,694 | 5.12 | 6.46 | 8,486 | 6.04 | 7.40 |
| DNA elements | 46,961 | 12.55 | 11.78 | 19,863 | 5.38 | 6.78 | 20,526 | 5.61 | 6.88 |
| Unclassified | 79,538 | 23.93 | 22.45 | 42,803 | 14.17 | 17.87 | 37,306 | 17.76 | 21.79 |
| Totals | 155,952 | 54.19 | 50.85 | 72,725 | 24.94 | 31.45 | 66,754 | 29.73 | 36.47 |
| Non-element repeats |  |  |  |  |  |  |  |  |  |
| Simple Repeats | 4,223 | 0.21 | 0.20 | 5,184 | 0.37 | 0.46 | 6,628 | 0.32 | 0.39 |
| Low Complexity | 2,833 | 0.18 | 0.17 | 4,377 | 0.28 | 0.36 | 4,194 | 0.26 | 0.32 |

\*Percent of contig bases.

Comparison of syntenic regions between *Pt* and *Pgt* highlights that the genome expansion in *Pt* is due to disperse integration of repetitive elements. In total, 4,319 orthologs are found between the two species in syntenic blocks of between 4 and 52 genes (Supplemental Material and Methods). However, the size of the syntenic blocks in *Pt* are 30.1% larger overall than in *Pgt*; syntenic regions cover 46.7 Mb of *Pt* and 35.9 Mb of *Pgt*. By contrast, regions of *Pst* are 71.2% the size of syntenic regions of *Pgt*, suggesting a compaction of *Pst*; however this analysis is impacted by the high percentage of gap in the *Pst* assembly, reducing the resolution of blocks that can be detected. Within the expanded regions of *Pt* and *Pst*, there are larger blocks of repetitive sequence interleaved between the orthologs (Figure 1), highlighting that the genome expansion appears due to disperse integration of repetitive elements.

**Figure 1.**
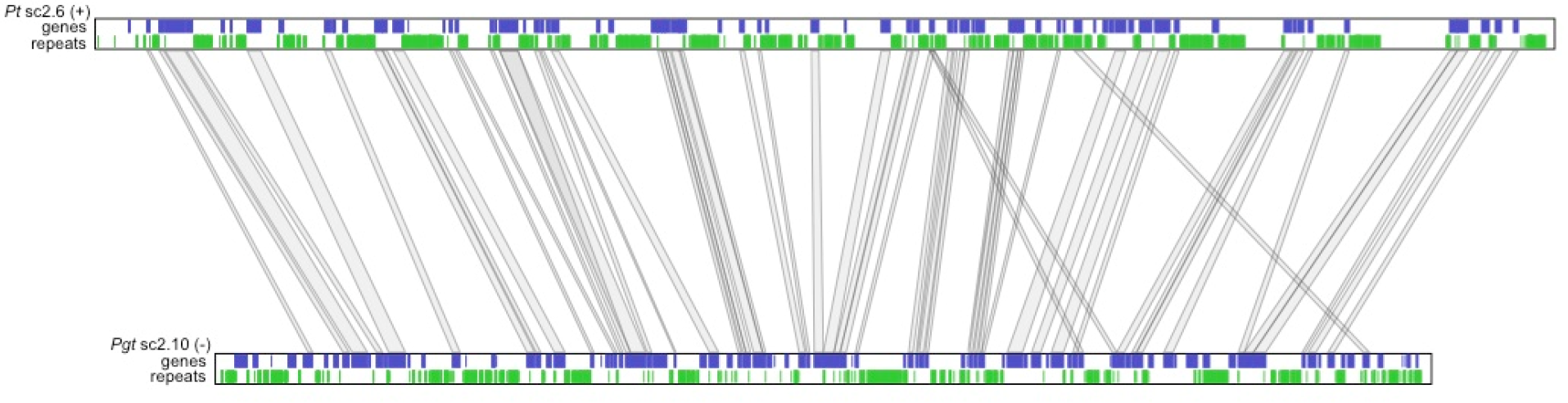
Genome expansion of leaf rust. Comparing syntenic regions of *Pt* and *Pgt* highlights the expansion of the *Pt* genome due repetitive element proliferation. The largest syntenic region between the two genomes is located on scaffold 6 of *Pt* and scaffold 10 of *Pgt*. Genes (blue) and repetitive sequences (green) are shown along each scaffold, with orthologs between the two genomes connected with lines. This region is expanded 1.34-fold in *Pt* relative to *Pgt*.

### High heterozygosity across all wheat rusts and evaluation of haploid assemblies

The cereal rust pathogens exist as dikaryotic (*n+n*) organisms for most of their life cycle, with a high level of heterozygosity between haplotypes. For *Pt*, 273,526 heterozygous single nucleotide polymorphisms (SNPs) were identified across the genome based on Illumina read alignment (Supplemental Material and Methods). Across the genome, the average rate of heterozygosity was 2.57 SNPs/kb, though a higher rate bias was observed in intergenic regions (2.85 SNPs/kb) than in genic regions (1.71 SNPs/kb). By contrast for *Pst*, 473,282 heterozygous SNPs were identified, for an average rate of 5.97 SNPs/kb; genic regions show a higher rate (7.49/kb) than intergenic regions (4.96/kb), a much higher rate than the 0.68 SNPs/kb previously reported for *Pst* with an assembly of the CY32 strain (Zheng *et al*. 2013). The rate of heterozygosity for *Pt* is similar to that reported for *Pgt* (Duplessis *et al*. 2011), where higher rates in genic regions (2.28 SNPs/kb) were found compared to intergenic regions (1.72 SNPs/kb), although the sequencing technology and methods differ between these studies. Heterozygosity levels in both *Pst, Pgt* and *Pt* are more than double those reported for genic and intergenic regions of *Mlp* (0.84 and 0.87 SNPs/kb respectively), supporting a high level of allelic diversity in these two wheat rust pathogens.

Regions of high heterozygosity could carry enough differences to prevent haploid assembly and could inflate the gene count for such regions, as alleles would appear as duplicated genes. Therefore, we examined the orthology assignment of genes found in all three wheat rust pathogens for conserved genes with two copies in only one species. Among the wheat rust fungi, *Pt* has only 230 species-specific two-copy paralogs (2:1:1; *Pt*:*Pgt*:*Pst*). *Pst* has an intermediate value of 361 species-specific paralogs while *Pgt* has 465 species-specific paralogs. Duplications detected by assembly self-alignment were most easily examined in *Pgt* due to the low scaffold count and only one region was found to have both haplotypes assembled separately (Duplessis *et al*. 2011). Overall this suggests that independent assembly of both haplotypes is minimal in all three wheat rust pathogens, as expected given the choice of assembly strategies that take heterozygosity into account.

### Core protein comparisons and orthology

To examine gene content variation between the three wheat rust pathogens and with other basidiomycetes, we compared the predicted proteins of *Pt, Pgt* and *Pst* to those of related genomes. These included *Mlp* and *Mli*, the smuts *Um* and *Microbotryum lychnidis-dioicae*, the fern parasite *Mixia osmundae*, the unicellular plant phylloplane “red” yeast *Sporobolomyces roseus*, the human facultative pathogen *Cryptococcus neoformans*, and the woodrotter *Coprinopsis cinerea*. By identifying orthologs across these genomes, we inferred the phylogenetic relationship of the species using single copy orthologs; *Pgt* and *Pt* are most closely related, with *Pst* being an earlier diverged outgroup (Figure 2), consistent with previous findings from phylogenetic analysis of the ITS ribosomal DNA region (Zambino and Szabo 1993). While *Pgt* and *Pt* are most closely related based on phylogeny, some features may be more conserved or have evolved in parallel in *Pgt* and *Pst*, which share the same alternative host.

**Figure 2.**
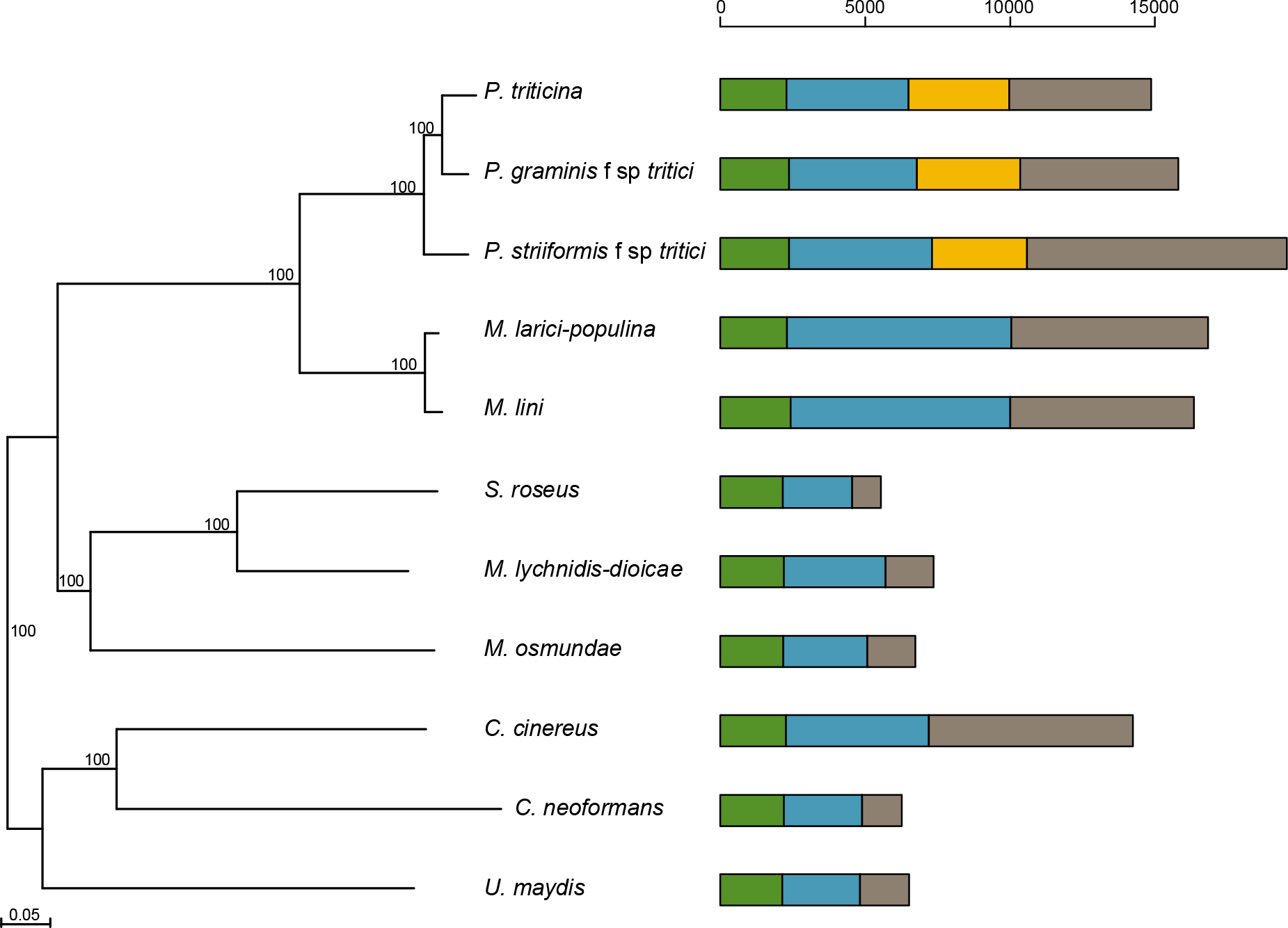
Phylogenetic relationship and gene conservation for 11 Basidomycete fungi. Orthologs were identified using ORTHO-MCL, and the aligned protein sequences of 1,208 single-copy genes were used to infer a phylogeny using RAxML. Bar graphs represent the core protein clusters shared amongst the species (green), conserved protein clusters (blue), and species specific protein clusters (grey). The orange bars represent protein clusters shared only among the wheat rusts. The horizontal scale indicates the numbers of orthologous groups.

The wheat rust pathogens have very different gene content from other Basidiomycetes, including a large fraction of species-specific genes. Less than half of genes in each wheat rust pathogen, an average of 6,867, were conserved among other basidiomycetes. All of the rust fungi (*Pt*, *Pst*, *Pgt*, *Mlp*, and *Mli*) contained an average of 6,482 species-specific predicted genes. Among the other compared basidiomycetes, only *C. cinerea* contained a similar number of species-specific genes. Among the wheat rust pathogens, *Pt* and *Pgt* had similar numbers of species-specific genes (5,443 and 4,901 respectively), while the *Pst* number was higher at 8,955. In addition, a large fraction of the genes are conserved across the wheat rust pathogens but not other fungi; an average of 3,440 are conserved in at least two genomes and 2,164 are found in all three.

To assess functional differences based on variation in genes between the three wheat rust pathogens and other fungi, we identified significant differences in the number of predicted protein domains. The three wheat rust pathogens were compared to *Mli, Mlp*, and to six other basidiomycetes to determine what protein families exhibit significant gain or loss in rust fungi. The majority of these are protein families involved with nucleotide binding and modification, transcription factor regulation, and protein modification (Table 3). These include the NAM-associated transacting factor family, the most significantly enriched protein domain overall; this domain is specific to the rusts among basidiomycetes, with 7 or 10 copies in *Mlp* and *Mli* and between 49 and 134 copies in the three wheat rusts. Three other Zn-finger transacting factor and a fungal-specific transcription factor families are also enriched. Four other enriched protein families are associated with carbohydrate active enzymes; trehalose phosphatase, pectinesterases, glycoside hydrolase family 26 (GH26), which processes mannan and galactomannan, and the GH76 family of alpha-1,6 mannanases. Other families are involved in carbohydrate processing and transportation, cell metabolism, and metabolite transportation (Table S3). A notable depleted family, NmrA, belongs to a family of transcriptional repressors involved in controlling nitrogen metabolite repression in fungi (Stammers *et al*. 2001). Other genes involved in nitrate metabolism are lost in *Pt* and *Pst*, as previously noted for *Pgt* (Duplessis *et al*. 2011). Overall these highlight recent adaptation of gene regulation and host-interaction via gene duplication and diversification.

**Table 3.** Significantly enriched protein families found in wheat rusts when compared to eight other species of basidiomycetes.

| Pfam Family | Enrichment ratio<sup>1</sup> | Q Value[1] Significance |
| --- | --- | --- |
| <u>Nucleotide-binding and modification</u> |  |  |
| PF14303.1 NAM: No apical meristem-associated C-terminal domain | 6.57 | 3.52E-100 |
| PF13873.1 Myb/SANT-like DNA-binding domain | 6.35 | 2.76E-04 |
| PF12776.2 Myb/SANT-like DNA-binding domain | 4.06 | 1.33E-28 |
| PF10443.4 RNA12 protein | 3.54 | 8.38E-03 |
| PF00891.13 O-methyltransferase | 3.52 | 6.43E-03 |
| PF13923.1 Zinc finger, C3HC4 type (RING finger) | 3.11 | 3.45E-31 |
| PF13639.1 Ring finger domain | 3.10 | 1.69E-38 |
| PF00097.20 Zinc finger, C3HC4 type (RING finger) | 2.88 | 2.44E-23 |
| PF13920.1 Zinc finger, C3HC4 type (RING finger) | 2.39 | 2.63E-03 |
| PF00270.24 DEAD/DEAH box helicase | 2.37 | 4.97E-19 |
| PF00271.26 Helicase conserved C-terminal domain | 2.09 | 3.04E-11 |
| <u>Transcription factor regulation</u> |  |  |
| PF10497.4 ZF-domain of monoamine-oxidase A repressor R1 | 4.63 | 1.71E-03 |
| PF00176.18 SNF2 family N-terminal domain | 2.56 | 5.77E-12 |
| <u>Cell damage defense</u> |  |  |
| PF00080.15 Copper/zinc superoxide dismutase (SODC) | 4.43 | 2.03E-11 |
| <u>Protein modification</u> |  |  |
| PF12861.2 Anaphase-promoting complex subunit 11 RING-H2 finger | 3.94 | 3.57E-16 |
| PF11793.3 FANCL C-terminal domain | 3.75 | 2.26E-02 |
| PF02338.14 OTU-like cysteine protease | 3.24 | 6.72E-06 |
| PF02816.13 Alpha-kinase family | 2.34 | 3.50E-02 |
| <u>Carbohydrate processing</u> |  |  |
| PF03663.9[Glyco hydro 76[Glycosyl hydrolase family 76] | 3.70 | 3.64E-03 |
| PF02156.10[Glyco hydro 26[Glycosyl hydrolase family 26] | 3.50 | 1.22E-02 |
| PF02358.11 Trehalose-phosphatase | 3.47 | 4.84E-05 |
| PF01095.14 Pectinesterase | 3.16 | 2.94E-02 |
| <u>Cell cycle control</u> |  |  |
| PF12678.2 RING-H2 zinc finger | 3.55 | 4.97E-19 |
| <u>Other</u> |  |  |
| PF03466.15 LysR substrate binding domain | 5.44 | 4.76E-02 |
| PF11937.3 Protein of unknown function (DUF3455) | 3.67 | 4.79E-02 |
| PF01637.13 Archaeal ATPase | 3.19 | 3.08E-03 |
| PF01735.13 Lysophospholipase catalytic domain | 2.75 | 1.60E-03 |
| PF13813.1 Membrane bound O-acyl transferase family | 2.57 | 3.32E-03 |

<sup>1</sup>log<sub>2</sub> weighted count wheat rusts/weighted count in non-wheat rust basidiomycetes

### Effector repertoire mining and conservation

Candidate secreted effector proteins (CSEPs) across the wheat rust pathogens are proteins predicted to be secreted and expressed during host infection are likely involved in host interactions. In this analysis, CSEPs predicted for *Pt* were compared to those previously identified in *Pst* and *Pgt*. From a starting set of 15,685 predicted proteins, including variant proteins encoded by alternate transcripts and novel genes detected by RNA-Seq data, a total of 1,358 CSEP-encoding genes were predicted for *Pt* (Figure S3). Of these, a total of 914 *Pt* CSEPs grouped in 385 families or ‘tribes’ previously assigned to *Pst* or *Pgt* CSEP tribes (Table S4; (Cantu *et al*. 2013)). From the remaining CSEPs, an additional 111 contained BLASTP sequence similarity (at ≤ e^−20^) to *Pst* and *Pgt* predicted proteins, of which 72 did not contain a predicted signal peptide at the expected initiation codon. The remaining 333 CSEPs were specific to *Pt* of which 246 were unique without any paralogs in the *Pt* protein set. The remaining 87 CSEPs belonged to *Pt*-specific gene families having from two to eight members per tribe. This highlights that while the vast majority of *Pt* CSEPs share sequence similarity with those in the other wheat rust pathogens, a subset are unique to each species. In addition, a disproportionate fraction of the wheat rust pathogen-specific genes mentioned above are predicted to code for secreted proteins. In *Pt*, a total of 17.0% of the wheat rust specific genes are predicted to encode secreted proteins compared to 9.6% of all predicted genes. This suggests the genes specific to the wheat rust fungi include a high fraction of potential effector proteins.

Based on gene ontology (GO) term assignment and similarities to Pfam domains, the molecular functions of the *Pt* CSEPs appear highly diverse, though some frequent categories were observed. The largest subcategories include a total of 123 CSEPs that have hydrolase activity, 76 contain ion-binding activity, and 44 have oxidoreductase activity (Figure 3). Pfam domain comparisons also revealed some potential common and unique functions among rust fungi (Figure S4). Since protein targeting is dependent on intrinsic protein motifs such as a nuclear localization signal or a chloroplast transit peptide, we analyzed all predicted CSEPs minus their signal peptides for their potential localization in the host to deduce possible functions. The distribution of their subcellular localization prediction in the plant indicates 388 CSEPs are potentially targeted to the cytoplasm, 361 to the nucleus, and 292 to plastids. A total of 190 of these proteins are targeted to membranes, 16 to the apoplast, seven to the Golgi system, four to the vacuole, and one to the ER (Table S4).

**Figure 3.**
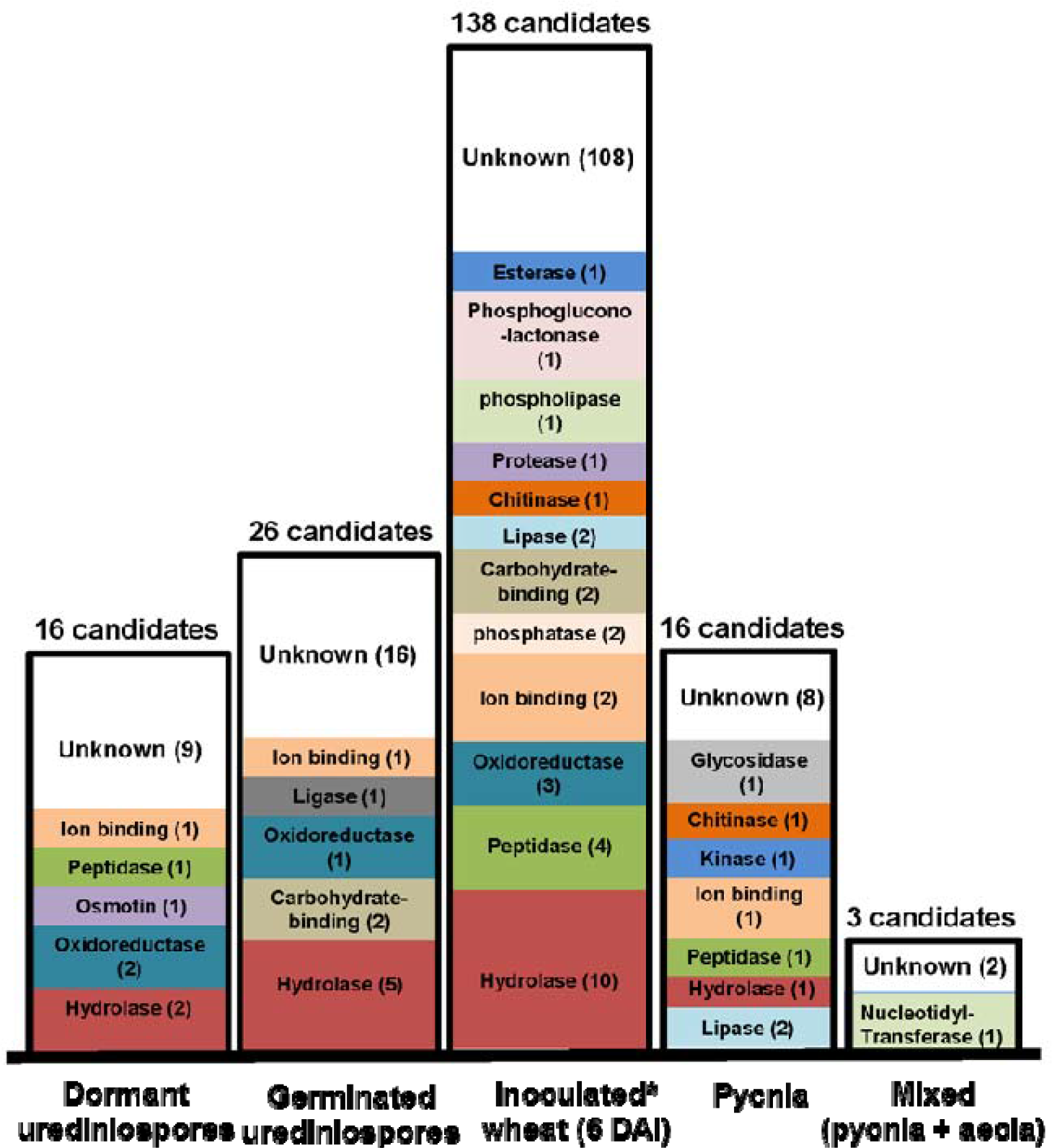
Highly expressed CSEPs vary across *Pt* developmental stages. The number of *Pt* CESPs with high transcript levels in each of the five RNA-Seq datasets compared to other datasets, with their predicted GO annotations. The fold change of the transcript levels in each two datasets was calculated. *Pt* effectors revealing a ≥ 5-fold change in one dataset when compared to the remaining datasets were selected and their molecular functions (GO annotation) were assessed. (*) *Pt* transcript level from inoculated wheat at (6 DPI) was calculated from an average of two samples derived
from independent wheat inoculations.

### Expression profiles across life cycle stages and two hosts

Functionally important gene expression was evaluated using RNA-Seq across diverse life cycle stages. RNA was sequenced from samples of dormant and germinating urediniospores, infected wheat leaf tissue at 6 DPI representing most of the infection structures of the uredinial spore genesis of the life cycle, and two stages collected from the alternate host, *Thalictrum speciosissimum* (meadow rue), the pycnia sexual stage and a later stage mixture of both pycnia and aecia. Comparing normalized counts across conditions revealed the two urediniospore samples were most highly correlated, and both were similar to the mixed sample of pycnia and aecia (Table S5). By contrast the pycnia and infected wheat leaf samples appeared the most different from the others.

To closely evaluate how secreted proteins change in expression across these conditions, all *Pt* CSEP genes were assessed and 199 effectors were identified with a minimum of 5-fold change. From these, 138 *Pt* effectors were highly induced in wheat-infected leaves (Figure 3) with 30 assigned to known proteins with hydrolase (10), peptidase (4), oxidoreductase (3), ion-binding (2), phosphatase (2), carbohydrate-binding (2), lipase (2), chitinase (1), protease (1), phospholipase (1), phosphogluconolactonase (1) and esterase (1) activities. Twenty-six CSEP genes were highly induced exclusively in germinating urediniospores. Among other stage-specific sets were 16 CSEP transcripts highly accumulated in dormant urediniospores compared to the germinated spore stage. When focusing on infection of the alternate host *Thalictrum*, 16 were highly expressed in pycnia and 3 during the later stage of the mixed pycnia and aecia sample (p+ae). In various datasets, except for the “mixed” pycnia + aecia stages, 1 to 10 candidate effector genes coding for proteins with hydrolase activity are highly induced. Among those, members of the glycoside hydrolases (GH) superfamily were prevalent, with GH18 specifically highly expressed (2 in wheat-infected tissues), GH16 and GH17 in germinating urediniospores, GH26 in dormant urediniospores, and GH16 in pycnia.

The predominant gene classes expressed in each stage were also examined by testing for functional enrichment in differentially expressed genes. Roughly half of the genes most induced during wheat infection compared to dormant spores encode predicted secreted proteins (Table S6). Other gene families that are enriched during wheat infection include the GH18 family, DNA binding proteins, peroxidases, and amino acid permeases (Table S7). By contrast, genes expressed in pycnia relative to dormant urediniospores are enriched for GMC oxidoreductase, LON proteases, potassium uptake, osmotic stress response, and chitin synthases (Table S8).

## Mating-type genes

### Pheromone receptors and precursors

Three homologs of the *Um* pheromone receptor Pra1 were found in each of the *Pt, Pgt* and *Pst* genomes (Table S9). All corresponding genes had a similar structure including five introns (Figure S5). The predicted proteins ranged in size from 379 to 395 amino acids, and had the characteristic seven transmembrane domains typical of these G-protein coupled membrane-inserted receptors (Bölker *et al*. 1992). A molecular phylogeny was calculated for these and the receptors from the poplar and pine rust pathogens, including known STE3/PRA proteins from several other basidiomycetes (Figure 4). This analysis revealed that the STE3 receptors from the Pucciniales formed a clade (blue boxes) well-separated from the Agaricomycotina (no color), the Ustilaginomycotina (orange box) and the Microbotryomycetes (yellow box). The rust clade encompassed two major groups, the subgroup with the STE3.1 homologs and an other branch where two subgroups represent likely more recent, lineage-specific duplication and divergence events (light blue and grey boxes).

**Figure 4.**
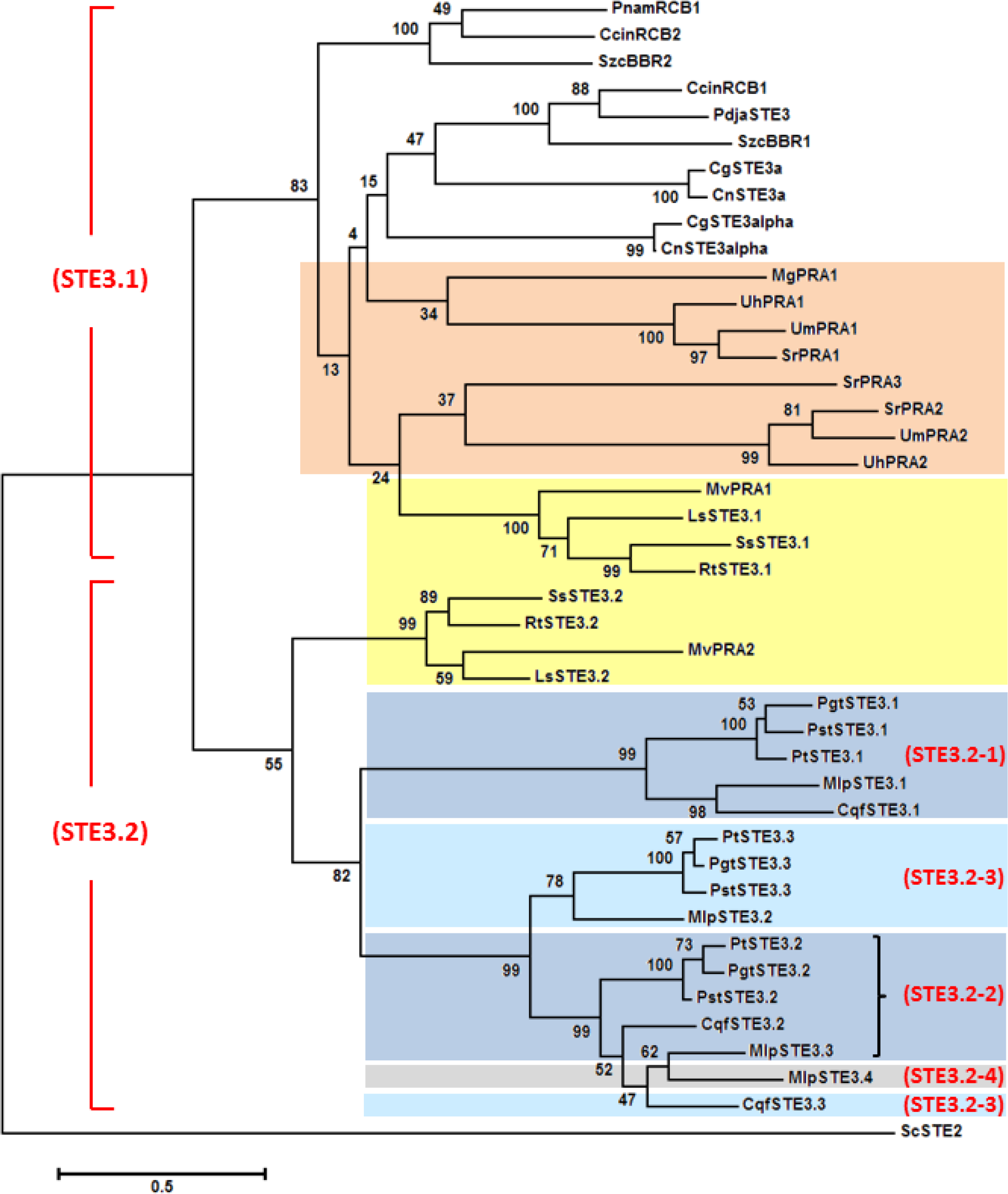
Molecular phylogenetic relationship of STE3-like pheromone receptor protein sequences of basidiomycetes. The pheromone receptor sequences were C-terminally truncated to exclude the cytoplasmic tail and to optimise the alignment (as in (Bakkeren *et al*. 2008); see Supplemental Materials and Methods for details), and were from *Coprinopsis cinerea* (CcinRCB1; CcinRCB2), *Cryptococcus gattii* (CgSTE3a; CgSTE3alpha), *C. neoformans* (CnSTE3a; CnSTE3alpha), *Leucosporidium scotii* (LsSTE3.1; LsSTE3.2), *Malassezia globosa* (MgPRA1), *Microbotryum violaceum* (MvPRA1; MvPRA2), *Pholiota nameko* (PnamRCB1), *Pleurotus djamor* (PdjaSTE3), *Puccinia* protein IDs are given in Table S9 (N-terminal 287 – 294 amino acids), *Rhodosporidium toruloides* (RtSTE3.1; RtSTE3.2), *Schizophyllum commune* (SzcBBR1; SzcBBR2), *Sporidiobolus salmonicolor* (SsSTE3.1; SsSTE3.2), *Sporisorium reilianum* (SrPRA1; SrPRA2; SrPRA3), *Ustilago hordei* (UhPRA1; UhPRA2), *U. maydis* (UmPRA1; UmPRA2). For the rust fungi: *Cronartium quercuum* f.sp. *fusiforme* (CqfSTE3.1; CqfSTE3.2; CqfSTE3.3), *Melampsora larici* f.sp. *populina* (MlpSTE3.1; MlpSTE3.2; MlpSTE3.3; MlpSTE3.4). STE2 of S. *cerevisiae* (ScSTE2) served as outgroup. Blue boxes: Pucciniales; orange box: Ustilaginomycotina; yellow box: Microbotryomycetes; Agaricomycotina, no color. STE names in red indicate tentative suggested groupings (see Discussion).

Using annotated *Pt* EST or known *mfa* sequences (Supplementary Note), three contigs in *Pt* and *Pgt* and two in *Pst* were found as containing putative *mfa* genes (Table S10). A putative *Ptmfa2* gene coding for a 33 amino acid protein with a characteristic C-terminal CAAX motif was identified on supercontig 2.517; while no gene model was initially predicted here, evidence of expression was detected in several life cycle stages (Figure S5). Extensive searches of genomic reads and RNA-Seq data could not identify other *mfa* genes. In all three *Puccinia* species, the predicted *mfa2* and *STE3.2* genes are divergently transcribed and are approximately 500-700 bp apart (605 bp in *Pt*; Figure S5), an organization reminiscent of *Ustilagomycete a2* loci. The *Pgt STE3.3* allele is 24 kb away from a potential *Pgt mfa* gene on supercontig 2.2 (Table S9, Table S10). In *Um* and *Sporisorium reilianum*, the *a2* loci each harbor two additional genes, *lga2* and *rga2* that are located in between the *Pra2* and *mfa2* genes. The LGA2 and RGA2 proteins localize to mitochondria and are implicated in mitochondrial fusion processes in that fungus (Bortfeld *et al*. 2004) but no obvious homologs could be identified in the *Puccinia* species.

### Homeodomain-containing transcription factors

Two allelic homologs of both HD1-and HD2––containing protein genes were found in each of the three *Puccinia* species that were termed *bE-HD2* and *bW-HD1* (Table S11). Gene models in the genome assemblies were found to be partial, and complete transcript sequences were constructed using *de novo* RNA-Seq assemblies (Figure S6). The predicted *Puccinia* HD2 proteins are approximately 374 amino acids in length whereas the HD1 proteins are approximately 620 amino acids in length. *PtbE2-HD2* and *PtbW2-HD1* are located close together and are divergently transcribed (Table S11; Figure S7). However, in the fragmented genome assembly, *PtbE1-HD2* and *PtbW1-HD1* are each located on a small contig so no direct inferences of linkage could be made for this pair. Comparative analysis of aligned DNA and protein sequences for the two alleles of each *PtbE* and *PtbW* gene revealed the conserved HD-specific domains within an overall conserved C-terminal region whereas the proteins were more diverged at the N-terminus, similar to the paradigm established in *Ustilago* species (Figure S6). A similar pattern of conservation was noted for the corrected *Pgt* and *Pst* alleles. A molecular phylogeny was generated to establish the relatedness among the HD-containing mating-type proteins in the three cereal rust fungi, compared to single homologs from the poplar and pine rust fungi (Figure 5). The allelic variants were closer to each other in each *Puccinia* species as they were among the species, since they are alleles and their sequences are evolving in a concerted fashion. Thus, among the rust fungi compared, the HD1-and HD2-containing transcription factors are each separated in defined clades, as is seen when many basidiomycetes are compared, indicating an ancient system in which allelic specificities are maintained because of their functionality (Bakkeren *et al*. 2008; Kues *et al*. 2011).

**Figure 5.**
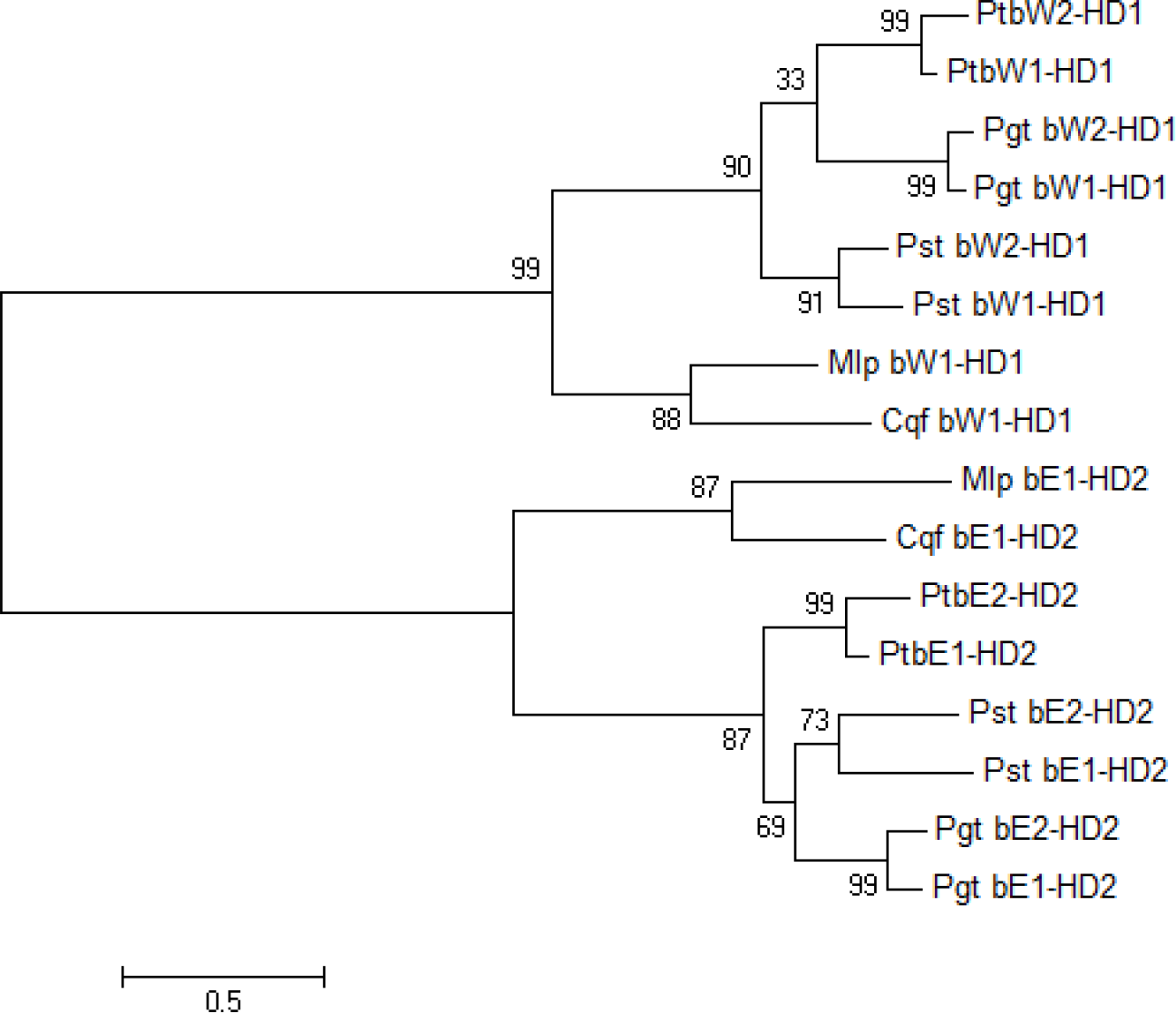
Unrooted tree displaying the molecular phylogenetic relationship of mating-type homeodomain-containing transcription factors in five rust fungi, using MEGA6
(Tamura *et al*. 2014). Maximum Likelihood method, showing percent bootstrap support (1,000 replicates) and branch lengths measured in the number of substitutions per site. The *Puccinia* allele sequences are given in Supplementary Figure S6. Cqf bE1-HD2, *C. quercuum* f.sp. *fusiforme*, jgi|Croqu1|661468; Cqf bW1-HD1, jgi|Croqu1|661465; Mlp bE1-HD2, *M. larici* f.sp. *populina*, jgi|Mellp1|124184; Mlp bW1-HD1, jgi|Mellp1|124183.

In all three wheat rust pathogens, a large contig with a complete divergently-transcribed pair of *bE* and *bW* genes is found while the other sometimes partial alleles are found on small contigs. This analysis highlights the challenges faced when assembling very similar sequences such as the conserved C-terminal domains, likely belonging to two different haplotype genomes. Therefore, to investigate the physical arrangement of both *bE-bW*pairs in the *Pt* race 1 genome, primers to the conserved 3’-ends of each gene (Table S12) were used in a PCR reaction, which yielded a single product of 3.9 kb from total gDNA isolated from germinating urediniospores. In dikaryotic urediniospores, both pairs are assumed to be present. Analysis of the sequences had revealed that nucleotide polymorphisms in restriction enzyme sites for Xma1 and Spe1 could be used to distinguish the allelic pairs. To verify whether or not the 3.9 kb PCR product contained both divergently-transcribed *bW*and *bE* gene pairs, it was digested with these enzymes for a prolonged period of time to indeed yield fragments consistent with the presence of both allelic pairs, confirming the suspected organization (Figure 6).

**Figure 6.**
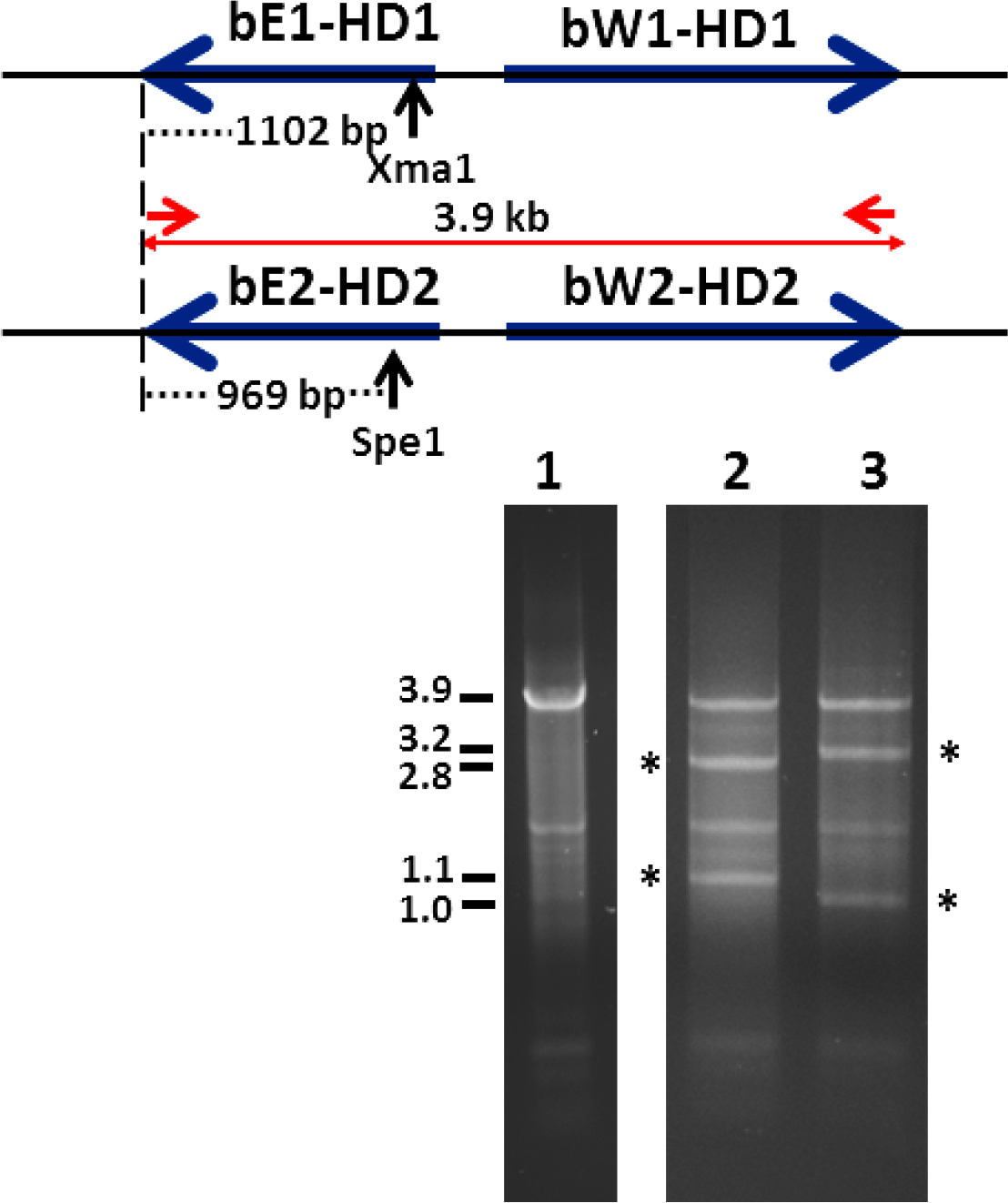
*Pt* has two complete *b* loci with divergently transcribed *PtbE* and *PtbW*genes. A 3.9 kb fragment was amplified by PCR from *Pt* gDNA from germinated dikaryotic urediniospores using conserved primers 1859 and 1902 (lane 1). Extensive digestion of this fragment by discriminating restriction enzymes Xma1 (lane 2) or Spe1 (lane 3) converted half of the PCR product to smaller fragments of the expected sizes in only either allelic pair (asterisks); the other allelic pair remained as a 3.9 kb PCR fragment.

### *Pt* HD genes are functional in *U. maydis*

We previously demonstrated the feasibility of using *Um* as a heterologous expression system for *Pt* genes (Hu *et al*. 2007). To examine the role of the candidate *Pt* mating-type genes, cDNA-derived *Pt* HD-containing transcripts were expressed in *Um*. Upon stable transformation of each of the *PtbE1-* or *PtbW1*-expressing constructs into either *Um* haploid strains *a1b1* or *a2b2*, the resulting transformants yielded cells that had changed morphology from growth by budding to a filamentous growth (Figure 7). When introduced into *Ustilago* cells, constructs expressing *b* mating-type genes of a different specificity or from different *Ustilago* species, transformants display these changed morphologies similar to regular mating interactions between cells of opposite mating types (Gillissen *et al*. 1992; Bakkeren and Kronstad 1993). This indicates that each of the PtbE-HD2 or PtbW-HD1 proteins are functional and can productively interact with the respective resident *Ustilago* b-gene subunit to initiate the switch to filamentous growth.

**Figure 7.**
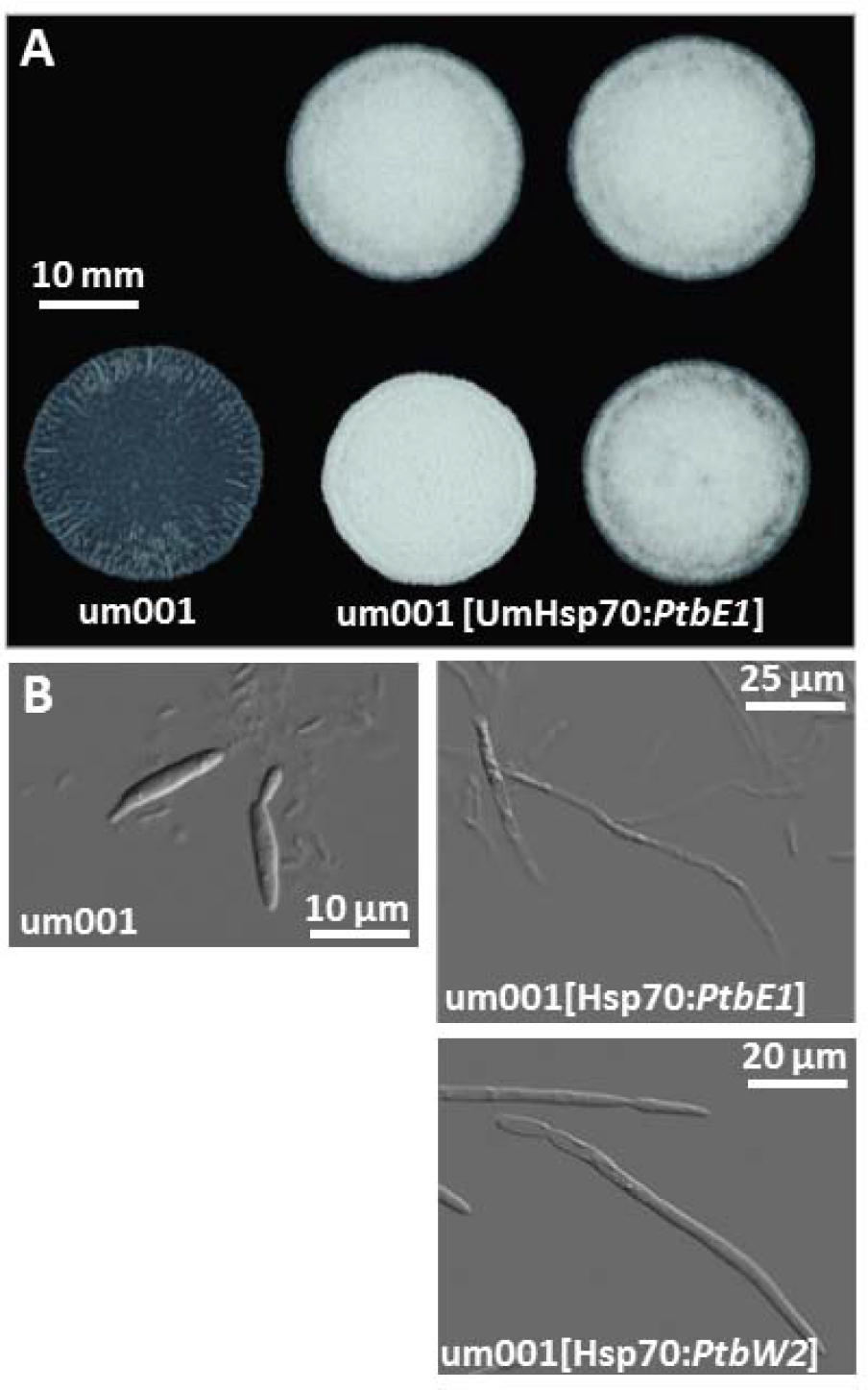
Single *Pt b* alleles are functional and interact with the *Um* b-regulated pathway controlling the switch from yeast-like to filamentous growth. In this representative example, *PtbEI* or *PtbW2* cDNA was expressed from the strong *Um* Hsp70 promoter and terminator elements. Four independent haploid *Um* integrative transformants displayed filamentous growth shown as aerial hypha when spotted on charcoal medium (Fuz+ colony phenotype; panel A) and by light microscopy (panel B).

Next we wanted to see whether or not a pair of Pt-specific HD-proteins could substitute for the resident pair in *Ustilago*. Two *Uh* strains of opposite mating type but each deleted for both *bE* and *bW* alleles (Uh553 (*a1 b0*) and Uh530 (a2 b0); (Bakkeren and Kronstad 1996) were each transformed with the above-tested single *Pt* HD-containing gene constructs (*PtbE1, PtbE2, PtbW1* or *PtbW2)*. Several independent stable transformants for each strain and construct were paired on a mating-type plate assay. Transformants of opposite mating type (*a1* vs. a2) should initiate proper cell fusion (brought about by the *pra* and *mfa* genes) allowing the respective *Pt* HD-proteins to interact; “Fuzz+” colony phenotypes would then be indicative of productive heterodimer formation and initiation of filamentous growth. Whereas control pairings of *Uh* Uh100 × Uh112 wild-type cells produced very “fuzzy” colonies after 48 hrs, all pairings of various combinations of transformants (5 per construct) did not produce colonies with significant aerial hyphae production. Upon microscopic analysis of the cells from such colonies, no convincing production of dikaryotic straight-growing hyphae as seen in wild-type mating interactions could be seen although mating hyphae were present and fusion initiated (data not shown).

### *Pt* mating-type genes are functional during wheat infection

Analysis of the transcriptomes revealed significant expression levels of almost all *PtSTE3* and *PtbE-HD2* and *PtbW-HD1* alleles during various life cycle stages, though expression in urediniospores was relatively low (Figures S8 and S9). Although expected to play a role during the sexual stage on the alternate host (certainly the pycniospore stage), a diversified role for them during infection could be envisaged. To examine such a role for the HD-containing alleles and *PtSTE3.1* and *PtSTE3.3*, the *Agrobacterium-mediated* HIGS technique was used (Panwar *et al*. 2013). Prior expression of silencing constructs in the wheat host targeted at these pathogen genes, significantly reduced fungal development as measured by biomass reduction and disease symptoms such as sporulation, upon infection with *Pt* urediniospores (Figure 8).

**Figure 8.**
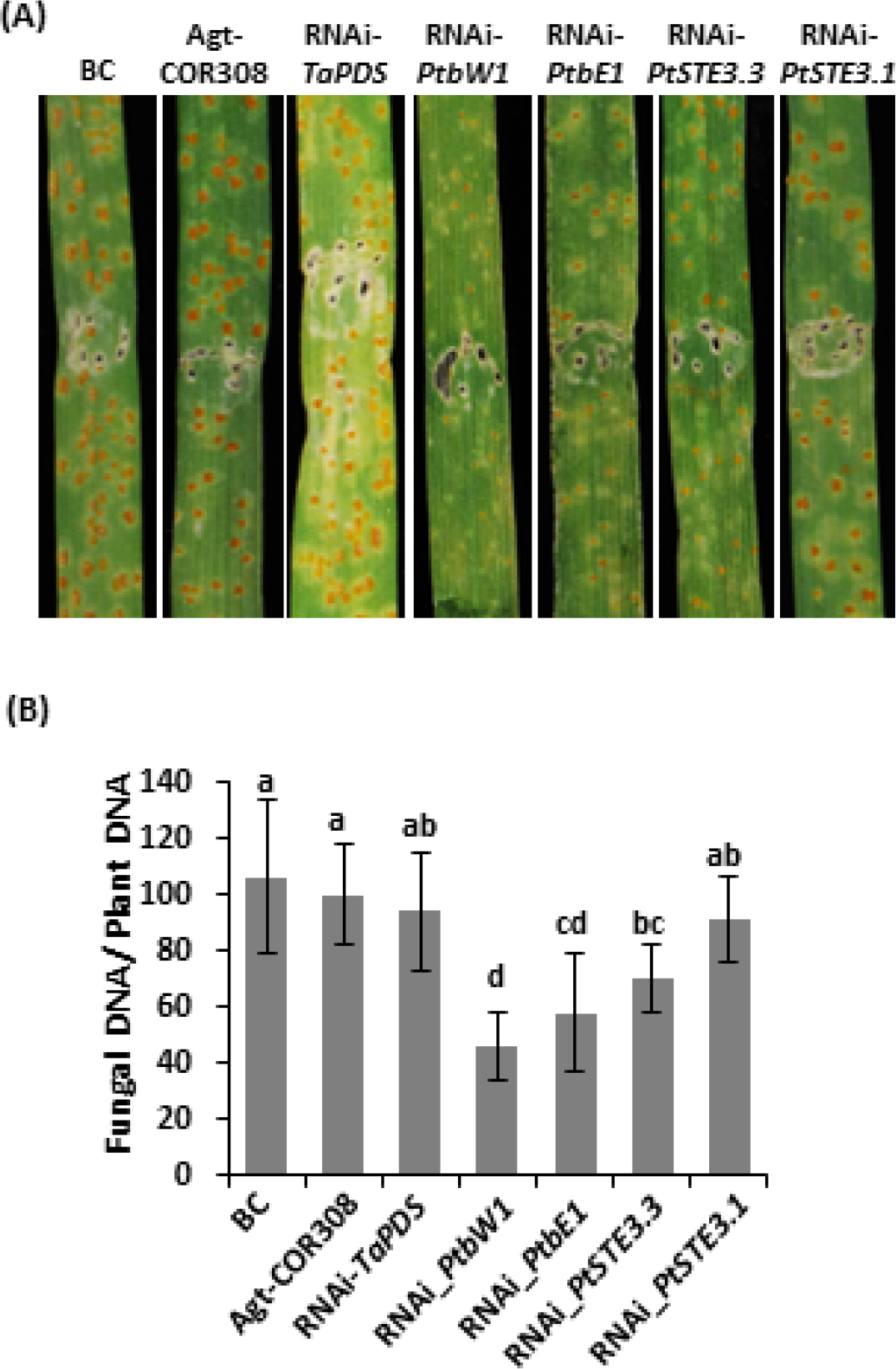
HIGS of *Pt* mating type genes by hpRNAi constructs. (A) Wheat leaf rust disease symptoms on wheat plants, infiltrated with *A. tumefaciens* delivering RNAi constructs and challenged four days later with *Pt* urediniospores, as compared with controls. Leaves photographed at 10 DPI. (B) Quantitative measurement of *Pt* DNA in infiltrated leaves: ratio of fungal nuclear to wheat nuclear genomes using single-copy genes *talicPtRTP1* (rust transferred protein 1) and wheat *TaEF1*, ±SD of three independent samples (pooled leaves) collected at 10 DPI. Different small letters above the columns indicate significance at *P* < 0.05 (Students t-test) as compared with Agt-COR308 control (*A. tumefaciens* without construct). BC, buffer control.

## DISCUSSION

The genomes of *Pt* and *Pst*, sequenced here and compared to those from *Pgt* and other Basidiomycetes, are notable for their expanded size and high level of heterozygosity. While each genome was assembled using different sequencing technology, each of the gene sets appears to be of similar quality, with high representation of core genes. The genome of *Pt* in particular has been expanded due to multiple classes of repetitive elements; while this higher repeat content was found to be dispersed across the genome assembly, repeat elements could impact the expression of nearby genes and could also contribute in this way to differences between related strains of the same species. Notably, we find that *Pst* has the highest level of heterozygosity and that this measure is larger than previously reported (Zheng *et al*. 2013). While some of this difference could be attributed to the isolate sequenced, the much larger size of the Pst-130 genome used in this previous study may result in an under-estimation of heterozygosity, such as in cases where both alleles of a gene were assembled independently.

Prior to this work, gene content surveys focused on genes expressed during infection and other life cycle stages. An extensive EST data set of 13,328 unique ESTs was created by sampling several stages in *Pt*, however, functional annotation was generally low (Hu *et al*. 2007; Xu *et al*. 2011). During this genome project, ESTs and newly generated RNA sequences were used to refine gene models and predict alternatively spliced forms in each of the genomes. Notably, *Pst* contained the largest set of predicted genes at 19,542, despite not having the largest genome. This total is similar to what has been found in other *Pst* genome projects. In the sequence of four other *Pst* races, the gene count varied from 18,149, to 21,030, which may have been impacted by differing levels of heterozygosity (Cantu *et al*. 2013). It is intriguing that in *Pst* there are many more CSEPs than in *Pt* or *Pgt*; in one study, 2,999 CSEPs were predicted in five consolidated *Pst* genomes, compared to 1,333 and 1,173 in *Pgt* and *Mlp*, respectively (Cantu et al., 2013). Virulence variability among *Pst* isolates is high and larger than for *Pgt* and *Pt*, likely due to a CSEP gene expansion and diversification to elude host recognition. In this regard, it may be significant that *Pst* can be found on 126 species of grasses among 20 genera (Line 2002; Cheng *et al*. 2016). Overall, the number of genes within the three rust fungal genomes is higher than that in other plant pathogenic fungi. Smut fungi have fairly low gene counts (6,500 – 7,000), but plant pathogenic fungi have as many as 17,735 in *Fusarium oxysporum* (Ma *et al*. 2010) and 16,448 in the necrotroph *Botrytis cinerea* (Amselem *et al*. 2011). *Mli* and *Mlp* have gene numbers of 16,271 and 16,399 respectively (Duplessis *et al*. 2011; Nemri *et al*. 2014), indicating a similar number of genes to wheat rust pathogens. Higher gene numbers may support the multiple spore stages and more complex life cycle in the rust fungi.

The large genome expansion in *Pt* due to repetitive elements was suggested by an earlier study of selected genome regions (Fellers *et al*. 2013). The genomes of other rust fungi are also enriched for repetitive elements though smaller in number and total DNA content. *Pst* and *Pgt* have similar repeat element numbers, while *Pt* is more like *Mli*, for which repeats occupy 87 Mbp or 46% of the genome (Nemri *et al*. 2014). While in some fungi the process of repeat-induced point mutation helps control the expansion of transposable elements, the activity of repeat-induced point mutation in the rust fungi (*Pgt* and *Mlp*) appears much lower than in other fungi (Amselem *et al*. 2015). Mobile elements are now considered to be essential “genome modifiers” that replicate and randomly re-insert to drive recombination, addition and/or deletion events, sometimes leading to protein neo-functionalizations. Regions of the genome enriched in repetitive elements have also been shown to be a source of genetic diversity, particularly within effector repertoires of pathogens for possible adaptation to their hosts (Haas *et al*. 2009; Raffaele and Kamoun 2012; Ali *et al*. 2014).

Similar to two previously sequenced wheat rust pathogen genomes (Duplessis *et al*. 2011; Zheng *et al*. 2013), 8% of the identified *Pt* transcript repertoire encodes potential secreted effectors. The three *Puccinia* species share a complement of secreted proteins, yet each has a group that is specific to its own species (Table S4 and Figure S4). Although all three are pathogens of wheat, their indigenous world-wide distribution and therefore evolutionary path and environmental (host) adaptation and life histories are different, as are their symptom formation and alternate host selection; this will have likely translated into a varied complement of CSEPs. Comparison among available rust fungus inventories allowed us to identify a preliminary set of CSEPs specific to the wheat rusts. However, poor annotation of candidate effectors, currently a common challenge in plant pathology, makes deducing biological meaning from specific subsets difficult. Nevertheless, based on Pfam domain searches, specific to wheat rust CSEPs were members of glycoside hydrolase families (GH15, GH17 and GH88), trehalose-phosphatases, members of the DyP-type peroxidase family, and glyoxal oxidase, and in addition, proteins with pro-kumamolisin, thaumatin and alcohol dehydrogenase-like domains (Figure S4). Intriguingly, 140 of the unidentified proteins were predicted to target the cytoplasm of the host and could be candidates with a role in the interplay with the host immune system.

Gene expression during the key stages of the fungal life cycle is quite different. Many CSEPs were strongly expressed in plant host tissues in comparison to the (germinating) urediniospore stages suggesting their particular role during infection. Although a large number of highly induced CSEPs could not be functionally annotated, a significant number fall into groups with hydrolase, peptidase and oxidoreductase activities. In the uredinial, pycnial, and aecial spore stages, many of the genes are associated with sugar, amino acid and membrane modification, or are amino acid transporters, nucleotide binding proteins or transcription factors. However, prior to uredinia formation, the fungus induces the protein manufacturing machinery and the most highly expressed genes are associated with ribosomes.

Mating and compatibility have been very difficult to study in the (cereal) rusts because many are macrocyclic, completing their sexual stage on a different (sometimes obscure or unknown) alternate host plant. Several studies have attempted to shed light on the mating-type system in rust fungi. Conclusions and speculations vary from rust fungi having a simple +/-bipolar system in several *Puccinia* and *Uromyces* species (Anikster and Eliam 1999) to a more complicated tetrapolar system with multiple allelic specificities in *Mli* (Lawrence 1980) and related oat crown rust pathogen, *P. coronata* (Narisawa *et al*. 1994). Our genome analysis demonstrates that the proposed simple +/− bipolar system is much more complex, at least in the cereal rust fungi. Analysis of the *STE3* genes in basidiomycetes supports the idea of two ancient clades, tentatively called STE3.1 and STE3.2 (in red in Figure 4). In members of the genus *Puccinia*, the STE3 pheromone receptors evolved in two major clades, which for consistency in this speculative scenario would then be called STE3.2-1 and STE3.2-2 (in red in Figure 4). Further duplication and divergence events likely led to the STE3.2-3 group, and an additional STE3.2-4 allele in *Mlp* (in red in Figure 4). Some of the genomic locations, synteny, and structure including the presence of multiple TEs and repeats we identified are in agreement with accelerated evolutionary potential in STE3-containing regions (Supplementary Note). This then would strengthen the idea that the bi-allelic recognition system is indeed ancestral in the basidiomycetes. In the Ustilaginaceae, S. *reilianum* has three *pra (STE3*) alleles (Schirawski *et al*. 2005) whereas in closely related *Uh* and *Um* only two are found although in the latter, one specificity is likely lost. A recent study among members of the Ustilaginaceae found three highly syntenic pheromone receptor alleles to be prevalent which led Kellner *et al*. (Kellner *et al*. 2011) to propose a tri-allelic recognition system to be ancestral. However, a recent study into the mating-type organization of a basal basidiomycete lineage, *Leucosporidium scotii*, strongly suggested a bi-allelic pheromone-receptor recognition system to be ancestral in the basidiomycetes (Maia *et al*. 2015). This is generally seen among the Ustilaginomycotina, the Agaricomycotina, and the Microbotryomycetes more recently identified. In the mushrooms, two clades of pheromone receptors are found but in each, expansion by duplication and mutation leading to many allelic series, is very common (Raudaskoski and Kothe 2010). Whether all three pheromone receptors in the *Puccinia* species are functional and capable of interacting with and responding to pheromones remains to be investigated. Since we analysed genome data derived from dikaryotic urediniospores from one isolate for each *Pt, Pst*, and *Pgt*, it is unclear which alleles represent opposite specificities normally expected for the pheromone response during mate partner interaction. Differing expression levels for specific alleles at different life cycle stages could indicate functional divergence and possibly a lost function in determining MAT-specificity such as seen in many mushrooms. Genome analysis of haploid-phase stages such as from several single pycnia on the alternate host, need to be performed to investigate this issue.

The limited number of *a* locus *Pra* and *mfa* alleles in smuts indicates a small repertoire of haploid fusion capabilities in nature (though promiscuity has been observed; Bakkeren and Kronstad 1996); this contrasts with multiple (allelic) arrays often found in mushrooms. Similarly, single *bE/bW* pairs are found in smuts with very few allelic variants identified in nature for the bipolars but many more (up to 33) for tetrapolar *Um*. The organization is often more complex in mushrooms where one to multiple HD1-HD2 pairs representing various alleles are found in arrays in many of their analysed genomes, accounting for the myriad of sexually productive specificities recognised in nature (Fraser *et al*. 2007; Bakkeren *et al*. 2008; Kues *et al*. 2011; Nieuwenhuis *et al*. 2013; Kües 2015). Closer to the Pucciniomycetes, in the Microbotryomycetes, a bipolar system with limited number of alleles for the HD-pair and *Pra-mfa* genes has been found in *Microbotryum violaceum* (Petit *et al*. 2012) but a “pseudo-bipolar” system with loose linkage of the HD-pair and pheromone-receptor genes, estimated to be 1.2 Mb apart, was described in *Sporidiobolus salmonicolor*, resulting in the discovery of multiple allelic HD-pairs in nature (Coelho *et al*. 2010). The *Puccinia* species genome analyses described here did not indicate close linkage of the *STE3/mfa* and HD genes (the current assembly and preliminary mapping data in *Pt* indicate these loci to be at least 216 kb apart; Supplementary Notes), though a loose linkage has not been ruled out. An inventory of HD alleles among a wide collection of isolates may answer some of these questions.

When introducing one particular *Pt bE* or *bW* allele into a wild-type haploid *Um* strain, these constructs triggered filamentous growth through the production of the respective HD protein, functional interaction with the *Ustilago* counterpart, and subsequent transcriptional activation of a subset of genes by the formed bi-species HD dimer, as shown for *Um* (Wahl *et al*. 2010). While we have shown such active interactions to occur between *b* alleles from different species within the smuts (Bakkeren and Kronstad 1993), such activity across quite diverged members of the basidiomycetes is astounding and suggests an ancient origin of these proteins indeed. However, the experiment introducing *PtbW1* and *PtbE2* alleles, each in a compatible *Uh* strain lacking *b* genes, did not trigger a switch to hyphal growth upon mating. Although fusion of mating hyphae was confirmed, this suggests that no productive interaction within the dikaryotic heterologous cell occurred. Given that we found only two allelic pairs of *bE* and *bW* in these *Puccinia* species and the overwhelming evidence of the productive interaction between such heterodimers in many very diverse basidiomycetes studied to date, it is unlikely that the PtbE and PtbW HD proteins would not interact. Failure to initiate filamentous growth in *Uh* then may indicate that the *Pt* HD-proteins lack domains or the specificity necessary for *Ustilago-specific* downstream interactions, nuclear import and/or for binding to *Ustilago* promoter elements that normally initiate the transcription of genes involved in the switch to filamentous growth (Scherer *et al*. 2006; Kahmann and Schirawski 2007); when *Pt-Uh* HD-heterodimers are formed, such functionality may be provided by the *Uh* component (Figure 7). Indeed, the predicted PtbE proteins are, at 374 amino acids, approximately 100 residues shorter than the *Ustilago* homologs. In addition, whereas the compositions of the helices that constitute the HD are relatively well-conserved between the *Pt* and *Ustilago* b-proteins, their location within the protein is significantly different. They may have evolved Puccinia lineage-specific adaptations.

The HIGS experiments demonstrated that some of the *Pt* mating-type genes were certainly functional in dikaryotic hyphae during wheat infection (Figure 8) in addition to the assumed activity during the sexual stage on *Thalictrum* spp. The involvement of the *a* mating-type genes not only in mating but also in pathogenicity of the dikaryotic cell type has been demonstrated in *Um* (Hartmann *et al*. 1996; Urban *et al*. 1996). Silencing of *PtSTE3.1* had less of an effect than of *PtSTE3.3* and this was correlated with the observed expression levels during wheat infection (Figure S8). As discussed above, since three STE3 genes are found, functional diversification catered to the various life cycle stages is distinctly possible. The sequences in the HD silencing constructs were designed such that they would silence both alleles and this was clearly detrimental to the infection process and shows therefore that they are important for pathogenicity. They could play a role in maintenance of the dikaryotic stage and/or induction or persistence of pathogenicity gene expression such as demonstrated for *Um* where the bE/bW heterodimer was shown to be essential for initiating the induction of a set of genes involved in the pathogenic life style (Brachmann *et al*. 2001; Wahl *et al*. 2010).

Wheat rust diseases are a major impediment to economic production of wheat in many areas in the world, and because of their rapid adaptation to newly introduced resistant cultivars and fungicides, they are a threat to envisaged increased yield for a growing population. Genome research on these elusive biotrophic pathogens has tremendously accelerated our understanding of their interaction with their host and the presentation of a *Pt* and another *Pst* genome and the comparative analysis to other rust fungi in this study has highlighted similarities and differences that can now be exploited for targeted crop protection strategies. Conserved and essential effectors, expressed during infection, and their intended host targets, would be interesting components for further study; a search for natural or engineered resistance genes recognizing such effectors could be effective.

## ACKNOWLEDGEMENTS

We thank the Broad Genomics Platform for generating DNA and RNA sequences, and the Michael Smith Genome Sciences Centre in Vancouver for sequencing the BAC ends. GB thanks M. Coelho for helpful discussions and acknowledges funding from the Canadian Genomics R&D Initiative. This project was supported by the USDA CSREES (awards 2008-35600-04693 and 2009-65109-05916).

*Mention of a trademark of a proprietary product does not constitute a guarantee of warranty of the product by the United States Department of Agriculture, and does not imply its approval to the exclusion of other products that may also be suitable. USDA is an equal opportunity provider and employer*.

## Author contributions

Designed the study and experiments: CAC, GB, LJS, SH, XC, JPF. Sample collection & preparation, generation of constructs: GB, XS, XA, FL, YA, MB, JZL, MW, CY, MB, LJS, JPF. Performed experiments: GB, VP. Performed assembly and annotation: JMG, SY, QZ. Analyzed the data: CAC, GB, HBK, DJ, RL, SS, MB, JPF. Wrote the paper: CAC, GB, JPF, HBK.

